# Recombinant protein delivery enables modulation of the phototransduction cascade in mouse retina

**DOI:** 10.1101/2022.11.10.516027

**Authors:** Sabrina Asteriti, Valerio Marino, Anna Avesani, Amedeo Biasi, Giuditta Dal Cortivo, Lorenzo Cangiano, Daniele Dell’Orco

## Abstract

Retinal dystrophies of genetic origin are often associated with mutations in the genes involved in the phototransduction cascade in photoreceptors, a paradigmatic signaling pathway mediated by G protein-coupled receptors. Photoreceptor viability is strictly dependent on the levels of the second messengers cGMP and Ca^2+^. Here we explored the possibility of modulating the phototransduction cascade in mouse rods using direct or liposome-mediated administration of a recombinant protein crucial for regulating the interplay of the second messengers in photoreceptor outer segments. The effects of administration of the free and liposome-encapsulated human guanylate cyclase-activating protein (GCAP1) were compared in biological systems of increasing complexity (*in cyto, ex vivo*, and *in vivo).* Analysis of protein biodistribution and direct measurement of functional alteration in rod photoresponses show that the exogenous GCAP1 protein is fully incorporated into the mouse retina and photoreceptor outer segments. Furthermore, only in the presence of a point mutation associated with cone-rod dystrophy in humans p.(E111V), protein delivery induces a disease-like electrophysiological phenotype, consistent with constitutive activation of the retinal guanylate cyclase. Our study demonstrates that both direct and liposome-mediated protein delivery are powerful tools for targeting signaling cascades in neuronal cells, which could be particularly important for the treatment of autosomal dominant genetic diseases.

## Introduction

The molecular processes underlying vision are triggered by the absorption of photons by opsins in retinal photoreceptors. Located in specific membranous compartments in the outer segments of rods and cones, opsins are G protein-coupled receptors (GPCRs) that activate the signaling cascade known as phototransduction. For many years, phototransduction has been considered paradigmatic for the largest class of GPCR-mediated signaling pathways (rhodopsin-like or class-A GPCRs), and the accumulated knowledge about the structural, biochemical, and physiological details of this cascade has enabled significant advances in drug design and pharmacological approaches for many other signaling pathways^1,2^.

The phototransduction cascade operates the conversion of the light-transported energy absorbed by the opsins into a chemical signal, i.e., the transient fall of vesicular glutamate release from the photoreceptor synaptic terminal, which is sensed by downstream neurons^3^. Rods and cones adapt to dramatic changes in ambient light by modifying the kinetics of phototransduction, a finely regulated process orchestrated by the second messengers Ca^2+^ and cyclic guanosine monophosphate (cGMP). Absorption of light by (rhod)opsin triggers the hydrolysis of cGMP by activating the phosphodiesterase 6, thereby causing the dissociation of cGMP from cyclic nucleotide-gated channels (CNG) and their closure. The ensuing decrease in the inflow of Na^+^ and Ca^2+^ hyperpolarizes the cell which, in turn, causes a reduction in neurotransmitter release. In parallel to these events, the light-independent extrusion of Ca^2+^ from the Na^+^/Ca^2+^, K^+^-exchanger leads to a drop of Ca^2+^ concentration in the outer segments (from ~600 nM in the dark to below 100 nM in bright light^4^).

These light-evoked alterations of second messenger levels in the photoreceptor outer segment trigger feedback mechanisms necessary for the timely shutoff of the cascade, as well as for the adaptation to specific light or dark conditions^3,5^. Subtle changes in Ca^2+^ concentration are promptly detected by guanylate cyclase-activating proteins (GCAPs), members of the neuronal calcium sensors family^6^. Two isoforms (GCAP1 and GCAP2) are expressed in rods and cones, but in human only GCAP1 seems to be actively involved in the phototransduction cascade as a modulator of retinal guanylate cyclase (GC) activity, the most prominent contribution arising from the GC1 isozyme^7^. In human photoreceptors GCAP2 is probably involved in biochemical processes other than phototransduction^8^, although its role in mouse phototransduction has been demonstrated^9^.

GCAP1, the main regulator of GC1, is a 23 kDa protein belonging to the EF-hand superfamily^10^ that ensures rapid detection of Ca^2+^ oscillations in the submicromolar range with a nanomolar affinity for Ca^2+ 11^. When Ca^2+^ concentration drops because of phototransduction activation, Ca^2+^ is replaced by Mg^2+^, which can bind in the same metal binding loops of motifs EF2, EF3 and EF4 (**Fig. 1a**)^12–14^ This mechanism allows GCAP1 to switch between different signaling states, namely Ca^2+^-bound (GC1-inhibitor) and Mg^2+^-bound (GC1-activator), regulated by specific allosteric mechanisms involving the protein, the metal cations and the myristoyl group bound at its N-terminal^14,15^. The conformation adopted by Mg^2+^-GCAP1 stimulates the synthesis of cGMP by GC1, thus permitting rapid restoration of dark-adapted cell conditions by reopening of the CNG channels^12,16^. The Ca^2+^- Mg^2+^ exchange results in relatively minor conformational changes for GCAP1^14,15^ (**Fig. 1a**), which are nevertheless sufficient to trigger the GC1 inhibitor-to-activator transition over the narrow physiological range of Ca^2+^ variation in the photoreceptor outer segment.

**Figure 1:**
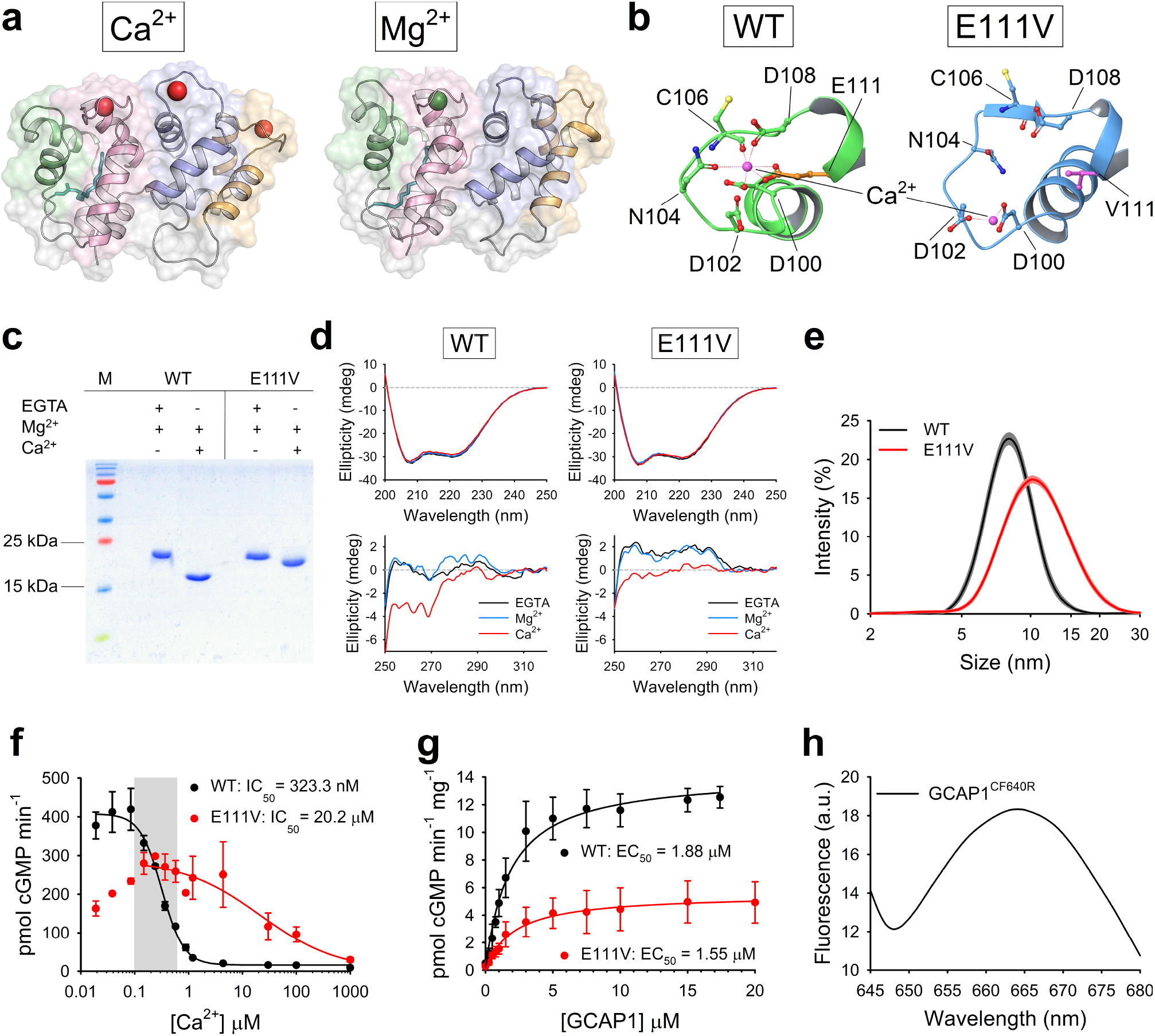
Biochemical and biophysical characterization of GCAP1 variants. **a)** Threedimensional structure of E111V-GCAP1 in its Ca^2+^-loaded (left) and Mg^2+^-bound (right) state after 200 ns Molecular Dynamics simulations. Protein structure is represented as cartoons with EF1, EF2, EF3 and EF4 colored in green, pink, blue and orange, respectively; the myristoyl moiety is shown as teal sticks, Ca^2+^ and Mg^2+^ ions are depicted as red and green spheres, respectively. **b)** Detail of the Ca^2+^-binding loop of EF3 in WT-GCAP1 (left) and E111V-GCAP1 (right) after 200 ns Molecular Dynamics simulations. Protein structure is shown as cartoons colored in green for WT-GCAP1 and blue for E111V-GCAP1; the sidechains of Ca^2+^-coordinating residues are labelled and are represented as sticks with O atoms in red, N atoms in blue, S atoms in yellow and C atoms in the same color as cartoons; the C atoms of E111 and V111 residues are colored in orange and magenta, respectively. Ca^2+^-ions are represented as pink spheres and labelled, zero-order bonds with Ca^2+^-coordinating residues are shown as dashed red lines. **c)** 15% SDS-PAGE of ~7 μM WT-GCAP1 and E111V-GCAP1 in the presence of 5 mM EGTA + 1 mM Mg^2+^ and 1 mM Mg^2+^ + 5 mM Ca^2+^. **d)** Representative far UV (upper panels) and near UV (lower panels) CD spectra of WT-GCAP1 (left panels) and E111V-GCAP1 (right panels) recorded at 25°C in PBS pH 7.4. Protein concentration for far and near UV was 10 and 33 μM, respectively. Far UV CD spectra were registered in the presence of 300 μM EGTA (black) and after serial additions of 1 mM Mg^2+^ (blue) and 600 μM Ca^2+^ (red), thus resulting in 300 μM free Ca^2+^. Near UV CD spectra were registered in the presence of 500 μM EGTA and after serial additions of 1 mM Mg^2+^ and 1 mM Ca^2+^ (500 μM free). **e)** Hydrodynamic diameter estimation by Dynamic Light Scattering of ~40 μM WT-GCAP1 (black) and E111V-GCAP1 (red) at 25°C in the presence of 1 mM Ca^2+^; standard errors are shown in grey and orange, respectively. **f)** GC1 enzymatic activity as a function of Ca^2+^ concentration (<19 nM – 1 mM) upon regulation by 5 μM WT-GCAP1 (black) or E111V-GCAP1 (red); cGMP synthesis was half maximal (IC_50_) at (323.3 ± 15.1) nM and (20.2 ± 7.6) μM with Hill coefficients of 2.16 and 0.99, respectively. The Ca^2+^-concentration range in photoreceptor cells is represented by the grey box. Data are presented as average ± standard deviation of 2 technical replicates. **g)** GC1 activity as a function of WT-GCAP1 (black) and E111V-GCAP1 (red) concentration (0-20 μM range); cGMP synthesis was half maximal (EC_50_) at (1.88 ± 0.15) μM and (1.55 ± 0.24) μM, respectively. Data are presented as average ± standard deviation of 3 technical replicates after normalization on the amount of GC1 present in cell pellets. **h)** Fluorescence emission spectrum of 5 μM GCAP1^CF640R^ upon excitation at 639 nm.

GCAP1 has been associated with autosomal dominant cone (COD) or cone-rod dystrophies (CORD)^17–31^, a class of severe inherited retinal diseases characterized by central vision loss, impaired color vision, and photophobia, resulting in photoreceptor degeneration^32^. Indeed, more than twenty point-mutations in the gene encoding GCAP1 (*GUCA1A*) have been found to be linked to COD or CORD. Recently, some of us identified a missense mutation in *GUCA1A* responsible for a particularly severe form of CORD. At the protein level, the mutation substitutes a glutamate residue in position 111 with a valine^24^. E111 is the twelfth residue of the Ca^2+^-binding loop of the EF3 motif and it is directly responsible for Ca^2+^-coordination by providing two negatively charged oxygen atoms from the carboxyl group (**Fig. 1b**); this bidentate ligation is fundamental to ensure the correct pentagonal bipyramidal geometry required for coordination of Ca^2+^-ions by seven oxygen atoms. The hydrophobic sidechain of V111, on the other hand, leads to a structural distortion of the EF3 loop, which becomes unable to coordinate Ca^2+^-ions (**Fig. 1b**), thus resulting in an 80-fold lower apparent affinity for Ca^2+^^24^.

COD and CORD remain incurable diseases, and the often autosomal nature of their transmission makes gene therapy-based approaches particularly challenging. In recent years, delivery of proteins and peptides to the eye have emerged as promising avenues for the treatment of a variety of ocular diseases^33^, although significant physiological and anatomical challenges remain^34^, especially when the goal is to modify biochemical processes occurring in the outer retina. In particular, to assist the development of effective therapies basic knowledge is needed on whether and how different proteins/peptides move across the ocular compartments^35^.

In this work, we explored the possibility of using direct or liposome-mediated administration of recombinant human GCAP1 to modulate the phototransduction cascade in mouse rods. The delivery experiments were first performed with a model eukaryotic cell culture, followed by an increase in complexity in which the biodistribution of proteins was assessed both *in vivo* and *ex vivo* in mouse retinas. Imaging experiments were complemented by functional ones, in which acute changes in flash responses were monitored while incubating the retinas *ex vivo.* The administration of the free and liposome-encapsulated protein were compared in each case. Our findings reveal that direct and liposome-mediated protein delivery are powerful tools for targeting signaling cascades in retinal neurons and could be particularly important for the treatment of autosomal dominant genetic diseases.

## Results

### Perturbed Ca^2+^-sensing properties of E111V-GCAP1 lead to constitutive activation of GC1

The purity and functionality of recombinantly expressed GCAP1 variants were verified by a combination of biophysical and biochemical techniques, to exclude potential effects of protein delivery treatments due to impurities or structural/functional defects. Ca^2+^-sensor proteins, including GCAP1, are known to modify their electrophoretic mobility in SDS-PAGE experiments under denaturing conditions^11^, depending on their Ca^2+^-loading state. Indeed, Ca^2+^-free proteins appear as single bands at their theoretical molecular weight, whereas their Ca^2+^-bound forms exhibit an electrophoretic shift to smaller apparent molecular weight proportional to their apparent Ca^2+^-affinity. We exploited this peculiar feature to assess both the purity of protein samples and the capability of WT-GCAP1 and E111V-GCAP1 to function as Ca^2+^-sensors (**Fig. 1c**). In the absence of Ca^2+^, both purified GCAP1 variants showed a single band around their theoretical molecular weight (23 kDa), which shifted to ~17 kDa in the case of WT-GCAP1 and to ~20 kDa in the case of E111V-GCAP1 upon Ca^2+^-binding, thus implying a substantial reduction in Ca^2+^-affinity for the pathological variant, confirming previous results from some of us^24^.

The structural response of GCAP1 variants to ion binding was monitored by Circular Dichroism (CD) spectroscopy, which allows monitoring changes in protein secondary and tertiary structure in solution at protein concentrations that mimic physiological ones (**Fig. 1d**). Both variants exhibited a far ultraviolet (UV) (200 - 250 nm) CD spectrum compatible with an all α-helix protein, with minima at 208 and 222 nm (**Fig. 1d**, top panels) and negligible variations in shape and intensity upon ion binding. This behavior was partly in contrast to that previously shown in ref.^24^, most likely attributable to the different buffer and temperature (PBS pH 7.4 and 25°C vs 20 mM Tris, 150 KCl, 1 mM DTT, pH 7.5 and 37°C). Concerning the tertiary structure (near UV CD spectrum, 250 - 320 nm), both variants displayed a significant rearrangement of aromatic residues upon Ca^2+^-binding in PBS pH 7.4 at 25°C (**Fig. 1d**, bottom panels), indicative of a change in protein tertiary structure. Mg^2+^-binding instead resulted in a minor conformational change, which was more pronounced in the case of WT-GCAP1. These results were substantially in line with the spectra recorded by some of us at 37°C, with minor differences attributable to the different buffer,^24^ which was found to affect also hydrodynamic diameter of GCAP1 variants (**Fig. 1e**). Both WT-GCAP1 and E111V-GCAP1 in PBS, pH 7.4 displayed a significantly larger hydrodynamic diameter ((8.68 ± 1.07) nm and (11.08 ± 0.07) nm, respectively), compared to their counterparts in 20 mM Tris, 150 KCl, 1 mM DTT, pH 7.5 buffer ((6.47 ± 0.03) nm and (6.08 ± 0.04) nm, respectively)^24^, as assessed by Dynamic Light Scattering. The slightly different experimental conditions of the present and the previous study^24^ lead to the same conclusions that the E111V substitution significantly impairs the Ca^2+^-sensitivity of GCAP1 with minor structural repercussions.

The enzymatic activity of the GCAP1-GC1 complex and its Ca^2+^-dependence creates a tight interconnection between Ca^2+^ and cGMP levels, which is crucial for both light adaptation and photoreceptor viability. Regulation of the GC1 enzymatic activity by GCAP1 variants was assessed both in terms of Ca^2+^ sensitivity and dependence on the level of protein regulator by measuring, respectively, the Ca^2+^ concentration at which GC1 activation is half-maximal (IC_50_) and the concentration of GCAP1 at which the synthesis of cGMP is half-maximal (EC_50_). Activation profile of GC1 by WT-GCAP1 exhibited an IC_50_ of (323.3 ± 15.1) nM (**Fig. 1f**), thus falling in the physiological intracellular Ca^2+^-range (<100 - 600 nM)^5^. On the other hand, the pathological variant E111V significantly dysregulated the activity of GC1, with an IC_50_ value ((20.2 ± 7.6) μM, p-value<0.05) ~63-fold higher than that of the WT, indicative of constitutive cGMP synthesis under physiological Ca^2+^ levels. Nevertheless, both variants displayed comparable EC_50_ values ((1.88 ± 0.14) μM for WT-GCAP1 and (1.55 ± 0.23) μM for E111V-GCAP1 (p >0.1), **Fig. 1g**), suggesting a similar affinity for the target enzyme, in line with previous results from some of us^24^.

### Liposome-mediated GCAP1 delivery to HEK293 cells

To assess the potential of liposome (LP)-mediated delivery of GCAP1 in biological systems of increasing complexity (*in cyto*, *ex vivo*, and *in vivo*) and investigate its biodistribution by minimizing the contribution of tissue auto-fluorescence (see Supplementary Materials), the far-red fluorescent dye CF640R was conjugated to the primary amines of solvent-exposed Lys residues of GCAP1 (namely, either of K8, K23, K24, K46, K87, K97, K142 or K162, **Movie S1**) to obtain the GCAP1^CF640R^ complex. SDS-PAGE confirmed the purity of protein samples and the success of the conjugation reaction (**Fig. S1a**). The effective removal of the unconjugated dye (**Fig. S1b**) and the number of CF640R molecules bound to each GCAP1 protein were then assessed by absorption spectroscopy (degree of labelling = 1.96, see Supplementary Materials). Finally, the emission fluorescence spectrum of GCAP1^CF640R^ (**Fig. 1h**) upon excitation at 639 nm was recorded before imaging experiments to verify the compatibility of the conjugated dye with our optical setups. The unconjugated dye (CF640R), GCAP1^CF640R^, WT-GCAP1 and E111V-GCAP1 were then encapsulated in LPs with a lipid composition corresponding to that of rod outer segment membranes (see Supplementary Materials for details).

The suitability of LP as carriers of small molecules and proteins was assessed by evaluating the size and monodispersion of the liposome suspensions loaded with different molecules. Regardless of the type of encapsulated molecule, Nanoparticle Tracking Analysis measured a LP diameter between (149.1 ± 3.0) nm and (168.7 ± 0.7) nm (**Fig. 2a, b**, and **c**, **Table S1**), with minor differences in both size and concentration (**Table S1**) up to 180 days (**Fig. 2d, e**, and **f**), suggesting that the LP suspension is stable over time. Finally, effective encapsulation of fluorescent molecules was also visually confirmed by the display of point-like fluorescence emission when LPs were filled with unconjugated CF640R and immobilized in agarose gel (**Fig. S2**).

**Figure 2:**
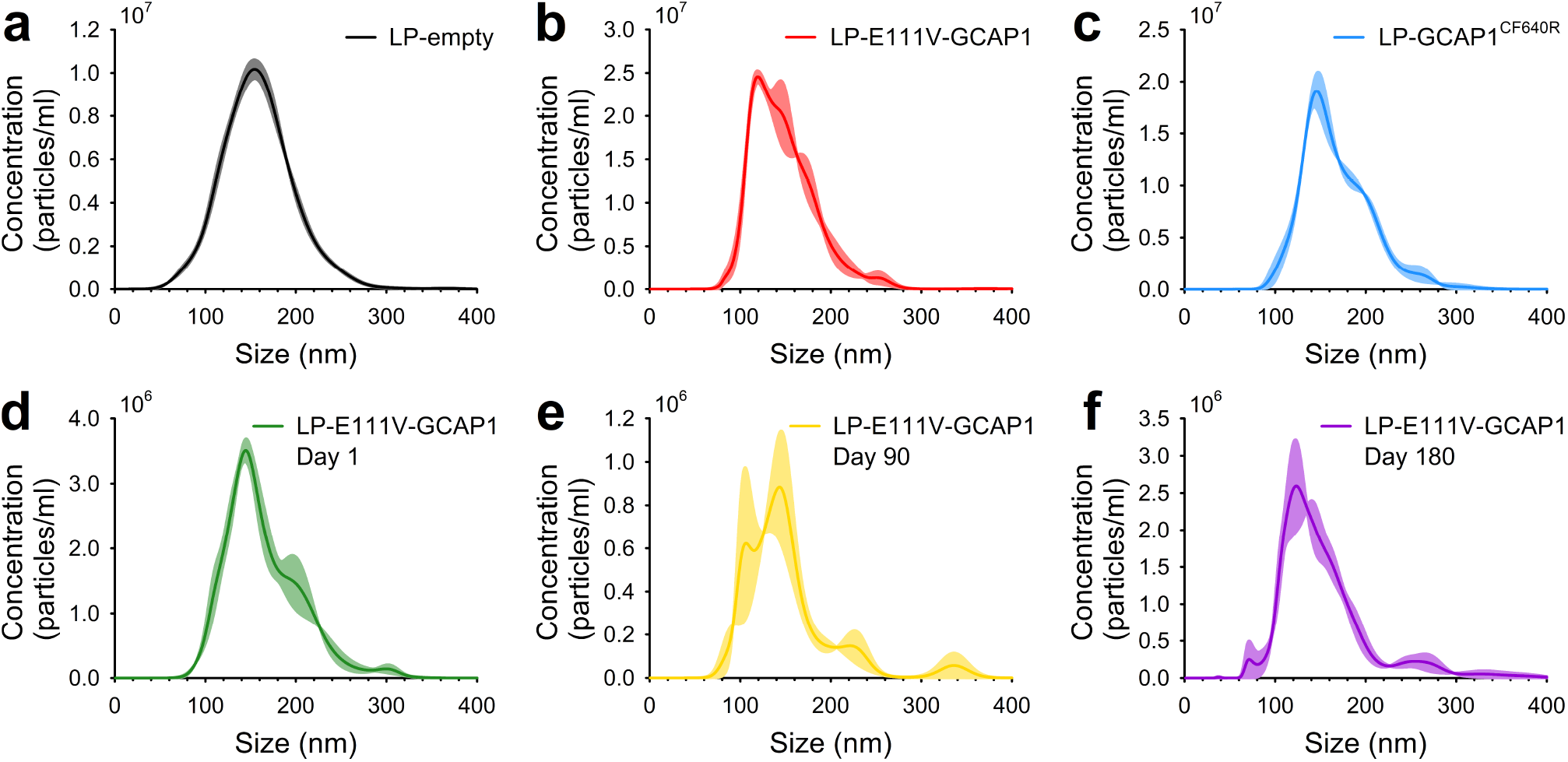
Nanoparticle tracking analysis of liposome preparations and their stability. Representative profiles of the size of **a)** 5.1 nM LP-empty (black), **b)** 4.3 nM LP-E111V-GCAP1 (red), **c)** 4.6 nM LP-GCAP1^CF640R^ (blue) estimated by Nanoparticle Tracking Analysis. Monitoring of the size of ~2.9 nM (**Table S1**) LP-E111V-GCAP1 after **d)** 1 (green), **e)** 90 (yellow) and **f)** 180 days (violet). Each plot represents the average of 3 independent measurements, standard errors are displayed as a lighter shade of the color of each trace. Concentrations refer to the stock solutions, before dilutions required for NTA analysis.

The capability of LPs to deliver recombinant GCAP1^CF640R^ was assessed on two different HEK293 stable cell lines. The first was transfected with pIRES plasmid encoding for eGFP and GC1 under separate promoters, characterized by a cytosolic green fluorescence (from now on cGFP); the second was transfected with pcDNA3.1 encoding for the eGFP-GC1 fusion protein, thus showing membrane green fluorescence (mGFP).

To address the potential direct membrane uptake of free-CF640R due to its small size (~832 Da) we monitored the cGFP cell line for 6 h (**Movie S2**) and visualized the mGFP cell line for 6 h and 24 h after incubation with free-CF640R (**Fig. 3a**). The absence of intracellular red fluorescence in both cell lines after 6h and 24 h suggested that free-CF640R is *per se* unable to penetrate cell membranes. Indeed, a diffused red fluorescence signal was observed only in the extracellular milieu, in the same time frames (**Fig. 3a**). This signal is likely attributable to the presence of phenol red in the medium as in real time imaging no medium replacement was performed.

**Figure 3:**
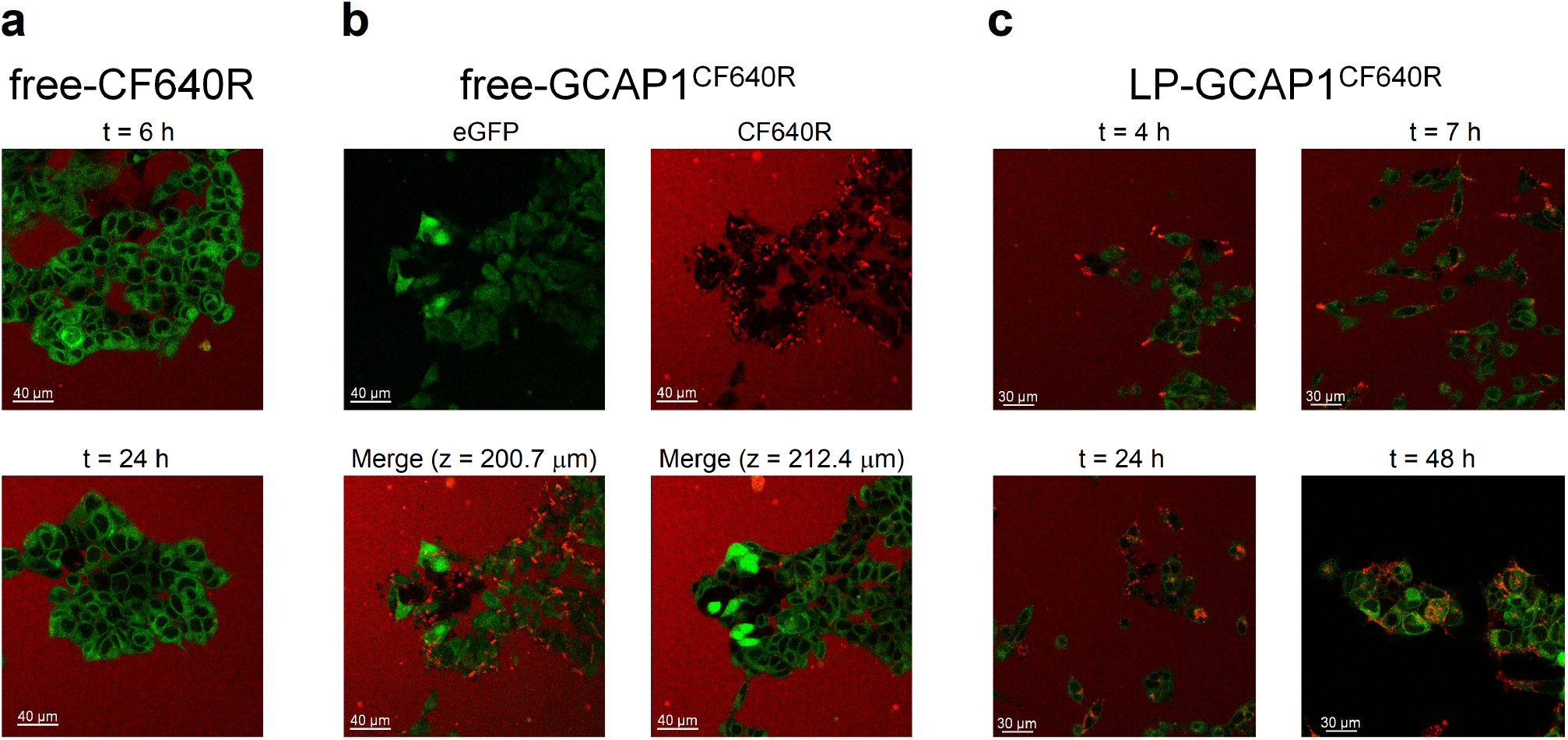
Live imaging of HEK293 cells incubated with free-CF640R, free-GCAP1^CF640R^ and GCAP1-encapsulating liposomes. **a)** representative images at a fixed z-plane of the mGFP cell line incubated with 100 μl of 140 μM free-CF640R after 6 h (top) and 24 h (bottom); **b)** representative images of the mGFP cell line after 24 h incubation with 100 μl of 104 μM free-GCAP1^CF640R^, top panels show eGFP (left) and CF640R (right) fluorescence, bottom panels show the merged fluorescence at z = 200.7 μm (left) and z = 212.4 μm (right). **c)** Live-cell imaging at the same z-plane of the mGFP cell line after 4 h, 7 h, 24 h, and 48 h incubation with 100 μl of 4.3 nM LP-GCAP1^CF640R^ (containing the same number of GCAP1^CF640R^ molecules in the aqueous core as compared to the free protein solution case). After 24 h the cell medium was replaced with FluoroBrite DMEM to avoid interference from phenol red, which gave rise to the red background fluorescence present in all but bottom right panel.

Similar experiments performed with fluorescently labelled GCAP1 (GCAP1^CF640R^), showed punctate fluorescence spots which tended to accumulate on the surface of mGFP cells after 6 h (**Movie S3**; punctuated red fluorescence is attributable to the interaction of GCAP1^CF640R^ with other molecules in the cell medium or with cell membrane, at odds with the diffused phenol red fluorescence). The intracellular space was not reached by the labelled protein even after 24 h (**Fig. 3b**, top panels). Further analysis at specific z-plane values (bottom panels in **Fig. 3b**) clearly confirm that the accumulation of fluorescence attributed to GCAP1^CF640R^ is limited to the cell membrane, as no signal was detected at the intracellular space. To confirm this finding, we repeated the same experiment with cGFP cells, which would allow the detection of overlapped green and red fluorescence signals in case of GCAP1^CF640R^ entering the intracellular milieu. Indeed, this was not observed in 6 h (**Movie S4**). To dampen the contribution of diffused red fluorescence from the extracellular milieu, we replaced the cell medium with FluoroBrite DMEM, which does not contain phenol. The same results were confirmed (**Fig. S3**), indicating that GCAP1^CF640R^ did not enter HEK293 cells in the observed timeframe.

As both cGFP and mGFP cell lines exhibited the same impermeability to free-CF640R and free-GCAP1^CF640R^, the capability of liposomes to deliver GCAP1^CF640R^ was tested only on the mGFP cell line. Live-cell imaging showed that 4 h after incubation (**Fig. 3c**, **Movie S5**), a punctuated red fluorescence, compatible with that emitted by LP-GCAP1^CF640R^, was accumulating on the cell surface. At t= 7 h, the same punctuated red fluorescence was distinctly detected in the cytosol, suggesting whole liposome internalization by the cells. Only after 24 h the fluorescence pattern started to change. While a more diffused red signal initially appeared at around 24 h, indicative of the release of GCAP1^CF640R^ from LPs, the observation of red and green colocalized fluorescence appeared to be completed at t=48 h (**Fig. 3c** and **Fig. S4**), suggesting an unhindered diffusion of GCAP1^CF640R^ in the cytosol. Finally, 48 h after incubation with LP-GCAP1^CF640R^ the cell medium was replaced with FluoroBrite DMEM to improve the signal-to-noise ratio, and a diffused colocalization of red and green fluorescence was clearly observed (**Fig. S4d**), thus confirming the complete intracellular release of GCAP1^CF640R^.

### Retinal distribution of GCAP1^CF640R^ and LP-GCAP1^CF640R^ following *ex vivo* incubation and intravitreal injection

#### Ex vivo incubations

To move to a higher level of biological complexity, we first assessed LP-mediated delivery of molecules by *ex vivo* incubation of isolated retinas. The rationale here was to eliminate variability related to *in vivo* transport across tissues, focusing solely on intraretinal mechanisms. Far red fluorescence was chosen because in preliminary tests we found that the extremely low tissue autofluorescence in this band greatly improved signal-to-noise ratio. Retina pairs (n=3) were incubated in parallel with 20 μl of LP-CF640R and LP-empty suspensions (at 3.9 nM and 5.1 nM, respectively) for 2 h at 37°C. After incubation retinas were rinsed and either viewed as wholemounts or slices, with the two partners placed adjacent in the dish. In all cases the LP-CF640R-treated retina showed fluorescence much above the control one. Unexpectedly, fluorescence was unevenly distributed across the thickness of the retina, being stronger in the inner layers (**Fig. 4a**). Nonetheless, even the outer retina showed some signal. Control slices had a flat autofluorescence profile, at the level of the chamber background. Interestingly, in the LP-CF640R-treated retinas many cell bodies in the ganglion cell (GC) and inner nuclear layer (INL) were markedly fluorescent. In the outer retina cones were occasionally clearly distinguishable (**Fig. 4b**), albeit dimmer than the aforementioned neurons. These data suggest enrichment of LPs in specific cell types, although it remained unclear whether they were internalized as intact nanovesicles or their fluorescent cargo released in the cell. To determine whether LP encapsulation affects the tissue access of large molecules, we incubated retina pairs (n=10) with 20 μl of 88.4 μM free-GCAP1^CF640R^ protein solution or 4.3 nM LP-GCAP1^CF640R^ suspension (containing the same number of GCAP1^CF640R^ molecules in the aqueous core as compared to the free protein solution case) for 3.5 h at 37°C, followed by slicing. Area fluorescence in the inner nuclear layer (INL) and photoreceptors (ONL+IS+OS) was separately measured and averaged across a random sample of slices from each retina. As we had observed qualitatively for LP-CF640R, fluorescence was higher in the INL both following incubation with LP-GCAP1^CF640R^ (153% SD 34%; p<0.01, n=10; paired Wilcoxon test) and free-GCAP1^CF640R^ (142% SD 35%; p<0.001, n=12) (**Fig. 4c**). Furthermore, treatment with LP-encapsulated protein led to significantly higher fluorescence compared to free protein in both the INL (153% SD 50%; p<0.05, n=10) and photoreceptors (129% SD 18%; p<0.01, n=10) (**Fig. 4c**). Many neurons in the ganglion cell layer and INL were distinctly stained after both types of incubations (**Fig. 4c**, images). It must be noted that, even in the photoreceptor layer, signal from retinas incubated with the free protein was above the level of tissue autofluorescence. This was confirmed by comparing incubation with free protein and PBS for 3.5 h at 37°C (n=2; 152% and 162%) (**Fig. 4d**). In summary, both LP-encapsulated and free protein rapidly penetrate the retina and appear to be taken up preferentially by some types of neurons. Moreover, despite both retinal faces being exposed to the incubation solution, fluorescent GCAP1 preferentially accumulates in the inner layers.

**Figure 4:**
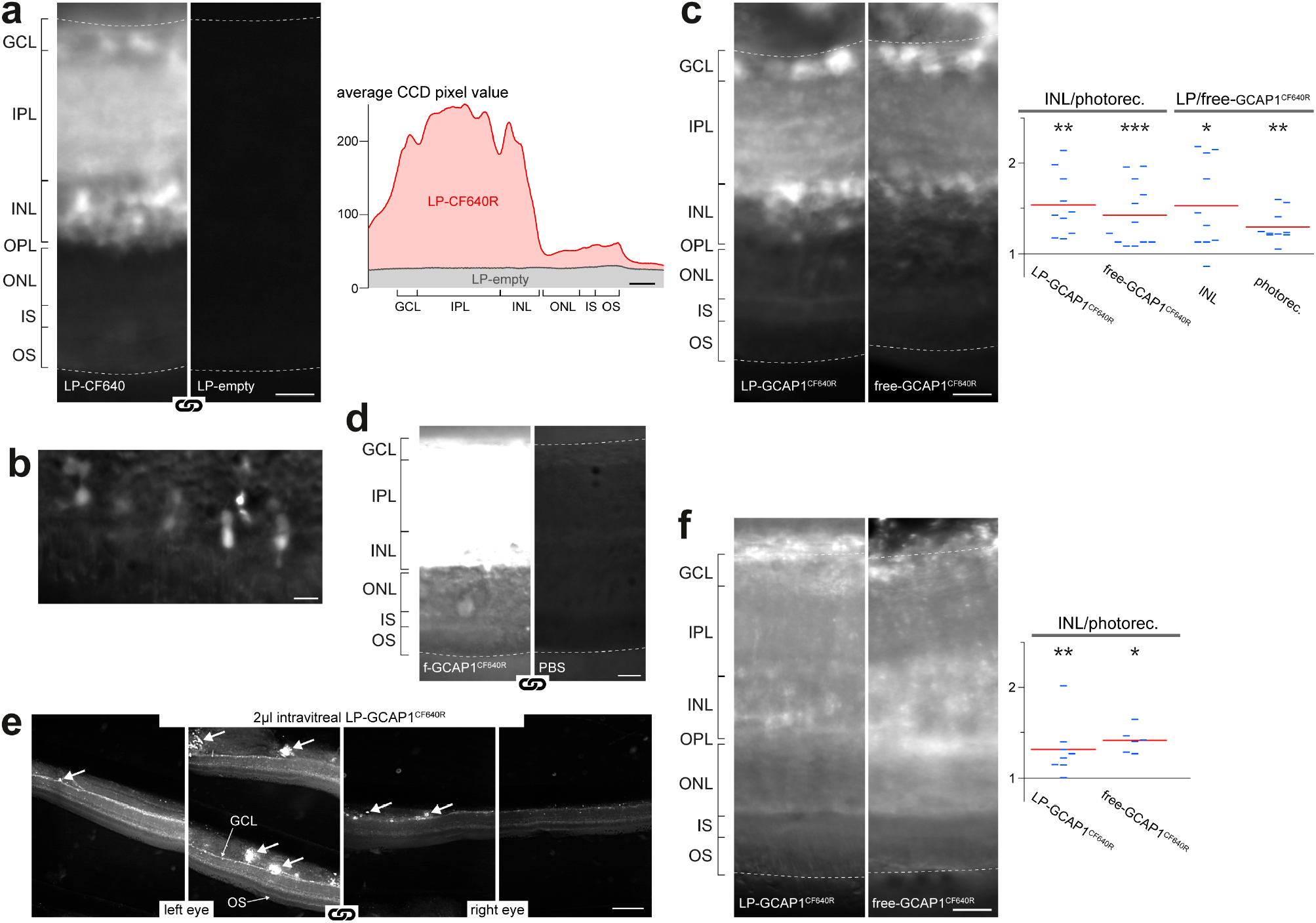
Biodistribution of free GCAP1 and liposome-encapsulated GCAP1 in mouse retinas following *ex vivo* incubation and *in vivo* intravitreal injections. **a)** Distribution of fluorescence in slices obtained from a pair of retinas incubated *ex vivo* with 20 μl of 3.9 nM LP-CF640R and 5.1 nM LP-empty, respectively: the same image acquisition and display parameters were used (chain link symbol). Plot shows the average fluorescence along the vertical axis of the same images. Scale bars 25 μm. **b)** Fluorescent cones in a slice from a retina incubated with 20 μl of 3.9 nM LP-CF640R. Scale bar 10 μm. **c)** Distribution of fluorescence after *ex vivo* incubation with 20 μl of 88.4 μM free-GCAP1^CF640R^ and 4.3 nM LP-GCAP1^CF640R^ (containing the same number of GCAP1^CF640R^ molecules in the aqueous core as compared to the free protein solution case). Scale bar 25 μm. Plots summarize data from several such experiments. INL/photorec: ratio of average fluorescence in the INL and photoreceptors layers (ONL+IS+OS), grouped by incubation solution; LP/free-GCAP1^CF640R^: ratio of average fluorescence after incubation with LP- and free-GCAP1^CF640R^, grouped by retinal layer (the same acquisition parameters were used in each retina pair). Blue dashes: data from single or pairs of retinas; red lines: means; *: p<0.05, **: p<0.01, ***: p<0.001 significantly different. **d)** Distribution of fluorescence after *ex vivo* incubation with 88.4 μM free-GCAP1^CF640R^ and PBS. The white point of the images was adjusted to enhance the outer retina: same acquisition and display parameters. **e)** Low magnification examples of retinal slices from the eyes of a mouse, both intravitreally injected with identical aliquots of 4.3 nM LP-GCAP1^CF640R^: the same acquisition and display parameters were used. Thick white arrows: zones of accumulation of fluorescence in the vitreous humor near the inner limiting membrane. Scale bar 250 μm. **f)** Examples of the distribution of fluorescence in retina slices after intravitreal injection with 2 μl of 88.4 μM free-GCAP1^CF640R^ and 4.3 nM LP-GCAP1^CF640R^ (containing the same number of GCAP1^CF640R^ molecules in the aqueous core as compared to the free protein solution case). Plots summarize data from several such experiments. INL/photorec: ratio of average fluorescence in the INL and photoreceptors in each retina, grouped according to injected solution. Scale bar 25 μm. In all images of this figure the focal plane lies deep in the slice.

#### Intravitreal injections

We went on to examine retinal delivery *in vivo* by injecting intravitreally 2 μl of 88.4 μM free-GCAP1^CF640R^ solution and 4.3 nM LP-GCAP1^CF640R^ (containing the same number of GCAP1^CF640R^ molecules in the aqueous core as compared to the free protein solution case) in the two eyes. After 20–24 h in darkness the animals were sacrificed, and their retinas isolated and sliced. We found that most injections led to some degree of retinal fluorescence (LP-GCAP1^CF640R^: n=9 out of 12; free-GCAP1^CF640R^: n=6 out of 11). Notably, while fluorescence was similar in different slices from the same retina, it varied greatly from eye to eye despite our utmost care in performing reproducible injections. This intrinsic variability was confirmed in a subset of animals in which both eyes were injected with identical solutions (n=3 pairs; **Fig. 4e**). Importantly, it hindered our ability to detect any significant differences in the delivery of LP-GCAP1^CF640R^ and free-GCAP1^CF640R^ to the retina (n=9 pairs). It is worth noting that residues of vitreous humor still adhering to the inner limiting membrane were often strongly fluorescent (**Fig. 4e** thick white arrows). In those retinas displaying fluorescence after intravitreal injection, the inner layers were significantly brighter than the photoreceptors, both in the case of LP-GCAP1^CF640R^ (132% SD 31%; p<0.01, n=8) and free-GCAP1^CF640R^ (142% SD 14%; p<0.05, n=6) (**Fig. 4f**). This mimicked what was observed in *ex vivo* incubations. However, following intravitreal injections individual cell bodies did not stand out above the overall fluorescence of the tissue (**Fig. 4f**): perhaps in these experiments there was sufficient time for uniform uptake by all neurons. In summary, we found that intravitreal injections are a viable, albeit rather inconsistent, means of delivery of free or encapsulated proteins to the retina.

### Delivery of E111V-GCAP1 induces a CORD-like phenotype in wild type mouse retinas

If a protein such as GCAP1 was able to gain access to retinal neurons in sufficient concentration, it could potentially be used to modulate biochemical processes^36^. As a proof of concept, we tested the functional effects on photoreceptors of E111V-GCAP1, known to be associated with CORD^24^. *Ex vivo* ERG recordings were made in a novel purpose-built chamber (**Fig. S5a**), which enabled prolonged incubation of retinas with relatively high concentrations of expensive test substances (i.e., using tiny overall amounts). ERG recordings were made at 35°C (except when stated otherwise) under pharmacological blockade of synaptic transmission to ON-bipolars (40 μM AP4)^37^. Except in one case, we did not remove the slow glial component with BaCl_2_ as in preliminary tests all effective concentrations also led to a progressive rundown of photoreceptor light sensitivity. The above conditions were associated to stable recordings of scotopic flash responses for over 4 hours. Two parameters were extracted: (*i*) light sensitivity measured by the flash intensity required to obtain a 50% response (i_50_); (*ii*) time to peak of the 50% response (TTP@i_50_) (see Supplementary Materials and **Fig. S5b**). These were normalized to their pre-treatment levels and processed to remove any trends also present in the control retina, leaving us with (ideally) the net effect of treatment (**Fig. S6**).

We first compared the incubation with 100 μl of 62 μM free-WT-GCAP1 to the same volume of PBS (n=14 animals; **Fig. 5a**). Over three hours of incubation no significant and systematic effects were detected either on sensitivity or kinetics (**Fig. 5a**). However, the incubation with 100 μl of 62 μM free-E111V-GCAP1 slowed response kinetics when compared to PBS (n=14; **Fig. 5b**), an effect highly significant already from the first minutes after delivery through the entire three hours of incubation (p<0.01). We confirmed this surprising result by comparing the same concentration (62 μM) of free-E111V-GCAP1 and free-WT-GCAP1, again observing a highly significant slowing of kinetics at most time points (n=8; **Fig. 5c**), which indicated that the effect is attributable solely to the E111V point mutation.

**Figure 5:**
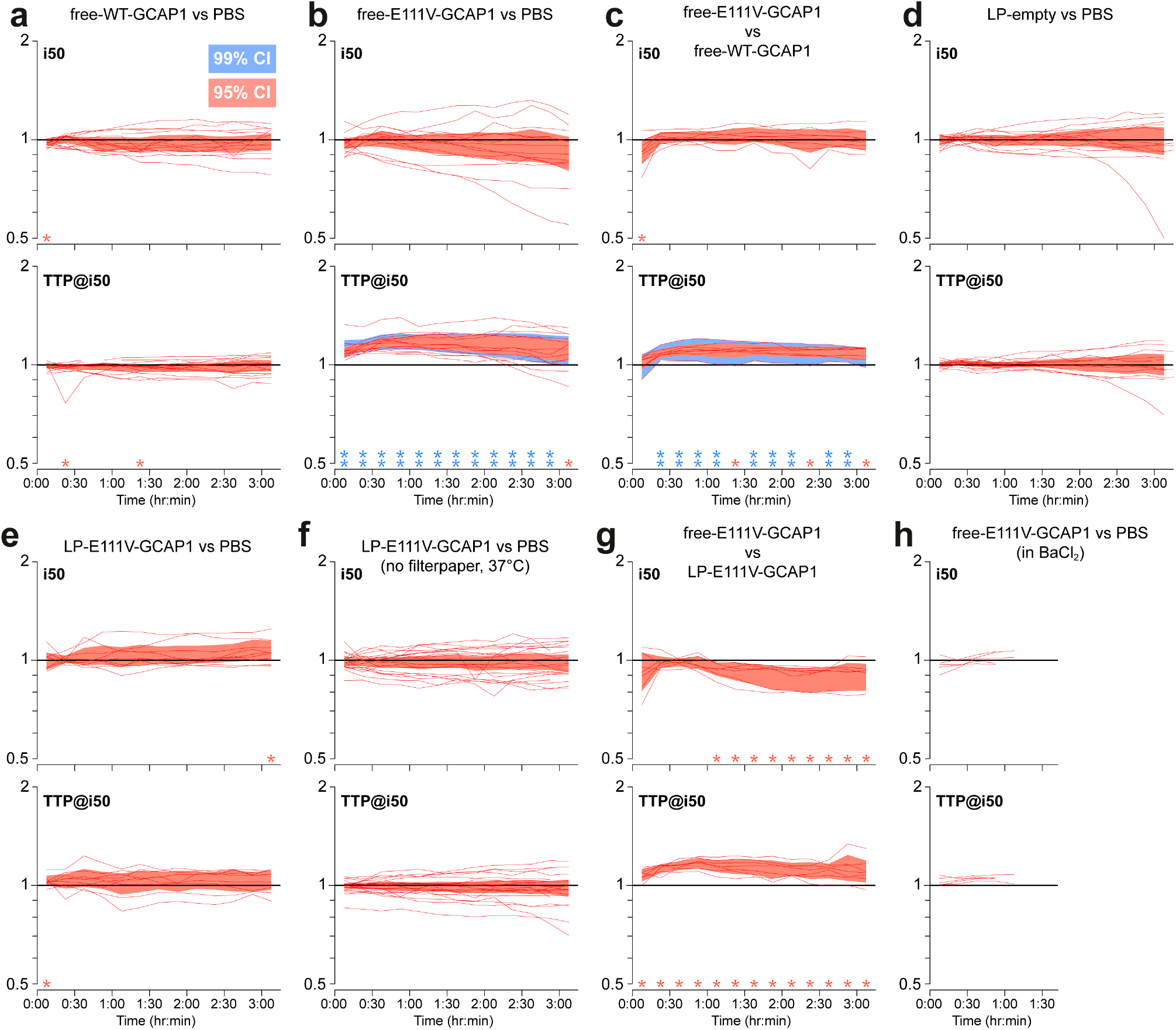
Functional effects on isolated retinas of incubation with free or LP-encapsulated recombinant GCAP1. The *ex vivo* ERGs of retina pairs were obtained in control conditions (Time < 0) and during parallel incubation with test and reference solutions for up to 3 h. Changes in light sensitivity (i_50_) and response kinetics (TTP@i_50_) were monitored by normalizing for pre-treatment control and reference solution (**Fig. S6**). **a-h**) Red lines represent individual experiments, each involving both retinas from an animal. Red (blue) shaded areas cover the 95% (99%) confidence interval. Loci above unity indicate a decrease in sensitivity or a slowing in kinetics. Red stars: p<0.05; double blue stars: p<0.01.

We then went on to examine the effect of LP encapsulation. When LPs-empty (5.1 nM) were compared to PBS no significant effects were detected (n=13; **Fig. 5d**), suggesting that LPs by themselves do not perturb phototransduction. More surprising was that LP-E111V-GCAP1 (4.3 nM, containing in the aqueous core the equivalent number of E111V-GCAP1 molecules present in a 62 μM solution) compared to PBS did not replicate the effects seen with the free mutant protein (n=9; **Fig. 5e**). Based on our experience with LPs holding fluorescent molecules we hypothesized that LPs might be scavenged by the filter paper supporting the retina in the chamber. To exclude this possibility, we modified our approach to hold the retinas in place during the recordings (**Fig. S5a**). Furthermore, to promote LP fusion/internalization in cells we raised the incubation temperature to 37°C. Despite these efforts no significant effects were detected over the course of three hours (n=21; **Fig. 5f**). We also compared the incubation of 100 μl of 62 μM free-E111V-GCAP1 with 100 μl of 4.3 nM LP-E111V-GCAP1 (containing the same number of E111V-GCAP1 molecules in the aqueous core as compared to the free protein solution case, n=7) and, given previous results, were not surprised to find a significant slowing of kinetics throughout incubation (**Fig. 5g**). Not seen in previous comparisons was a weakly significant increase in light sensitivity. Lastly, we confirmed that 100 μl of 62 μM free-E111V-GCAP1 slow response kinetics also when 50 μM BaCl_2_ is present in the bath solution (n=5; **Fig. 5h**), although these incubations had to be terminated after about 1 h for reasons stated above.

## Discussion

The *in vitro* characterization of GCAP1 variants used in delivery experiments showed that E111V-GCAP1 constitutively activates GC1 as a consequence of its impaired Ca^2+^ sensing, while ruling out the possibility that the effect is due to protein misfolding or altered affinity for the target. Furthermore, the GC1-GCAP1 system reconstituted in vitro showed a Ca^2+^ sensitivity perfectly in line with the intracellular Ca^2+^ changes that occur in photoreceptors during phototransduction activation, thus demonstrating the functionality of recombinant proteins.

Experiments with two different lines of HEK293 cells expressing GC1 clearly showed that fluorescently labelled GCAP1 (GCAP1^CF640R^) tended to accumulate near the membrane but did not cross it. In contrast, 4 h after incubation, the same liposome-encapsulated protein (LP-GCAP1^CF640R^) started to enter the cell and was clearly observed in the cytoplasm 24 and even 48 hours later.

Considering that HEK293 cells were impermeable to the unconjugated dye and that GCAP1^CF640R^ failed to cross cell membrane in a 24 h timeframe, this suggests that liposomes are indeed required to transport GCAP1 intracellularly in these cells.

A completely different scenario was observed in experiments performed with mouse retinas. Both free and LP-encapsulated GCAP1^CF640R^ were found to enter retinal neurons in the short time span of our *ex vivo* incubations (3.5 h). This also occurred 20–24 h after intravitreal injection with, however, a key difference: while in the former experiments specific neuronal cell bodies were highly fluorescent, in the latter ones fluorescence was locally uniform in the tissue. Perhaps the longer time frame of intravitreal injections compensates for varying rates of uptake by different neurons, thereby equalizing fluorescence. What the two techniques had in common was a preferential, but not exclusive, delivery of protein/LPs to the inner retina. This suggests that ocular diseases affecting retinal ganglion cells may be particularly well suited for protein therapy approaches relying on the delivery of recombinant proteins, either with or without the use of LPs as vectors.

Taken together, these fluorescence data suggest that cell membrane composition is important for the fate of free extracellular GCAP1. The lipid composition of HEK293 membranes^38^ significantly differs from that of photoreceptors, which is known to change during retinal development^39^ as well as between cone- or rod-dominant retinas^40^, and in pathological conditions^39,41^. The complexity of retinal lipid composition and metabolism could partly explain the differences observed in the two cell types. Indeed, liposomes with a lipid composition that mimics the photoreceptor membrane are apparently able to enter both HEK293 and retinal neurons, although the process takes at least 24 h in the former. However, a very surprising result, for which we currently have no explanation, is how exogenous GCAP1 can, in the absence of lipid carrier, cross retinal cell membranes and quickly achieve a relatively homogeneous distribution, at odds with HEK293 cells, where no protein internalization was observed even after 24 h. Moreover, the mechanism by which a homogeneous intracellular distribution of exogenous GCAP1 can be observed across different neuronal layers remains currently unknown. Perhaps the protein distribution among photoreceptors is somehow related to the recently discovered nanotube-like connections^42,43^ that allow the exchange of intracellular material^44^ including whole proteins, while the inner-to-outer retina protein exchange could be mediated by glial transcytosis operated by Mueller cells^45^. These hypotheses, which have tremendous implications for protein targeted therapy of retinal diseases, need further investigation. In a comprehensive set of electrophysiological experiments, we found that free human E111V-GCAP1 rapidly induces a significant slowing of the photoresponse, as measured by the TTP@i_50_. Crucially, the wild type protein did not evoke this effect, thereby pointing to a key role of the E111V point mutation. On the broader level this finding provides strong independent confirmation that free GCAP1 is taken up by retinal neurons. On the specific level of phototransduction it is striking, considering the compensating effect played by GCAP2 in murine photoreceptors^9,36,46^. In stark contrast, when the mutant protein was delivered encapsulated in LPs no such effects on kinetics were observed in the 3 h of incubation and recording. Considering the strong evidence that LP-encapsulated fluorescent GCAP1 ends up in retinal neurons relatively quickly, the functional data suggests that liposomes could be initially internalized intact, while only over the course of many hours release their cargo.

In a previous study we incubated mouse retinas for 2 h at 37°C with LPs containing either the homologous protein recoverin or an antibody against the same protein^47^. In that case, we observed a significant difference in saturating response kinetics between the two treatments. The lipid composition of those liposomes was somewhat different (phosphatidylcholine: cholesterol at various molar ratios) than that used in the present work. Also different was the electrophysiological recording technique (loose seal patch clamp from single rods), which did not require a pharmacological blockade of synaptic transmission or involve the presence of a slow glial ERG response component. Aside from these relatively minor differences, a key factor could be the relative ability of the different proteins of perturbing phototransduction. If, as postulated above, liposomes slowly release their cargo once inside the photoreceptors, a functional effect after 2-3 h may only be detectable when delivering a strongly impacting protein. Recoverin antibody could, in principle, possess such an effect, considering the crucial role of the recoverin-mediated Ca^2+^-feedback on rhodopsin kinase in accelerating the shutoff^48^. In contrast, in the present study the recombinant GCAP1 mutant had to outcompete the endogenous wild type protein

An unexpected finding of this study, highly relevant for retinal disease therapy, is that intravitreal injections show extreme trial-to-trial variability in the translocation of delivered molecules to the retina. Given our care in performing reproducible ocular injections we would rather attribute this phenomenon to complex flow dynamics or inhomogeneities in the vitreous. Whatever the mechanism, studies employing intravitreal injections should carefully consider whether variability in their observed therapeutic effects may have the same origin. Clearly, demonstrating significant effects of a drug candidate (or conversely excluding any medically relevant effects) may require a high number of test subjects.

Our study shows that direct and liposome-mediated protein delivery are powerful tools for targeting signaling cascades in neuronal cells and could be particularly important for the treatment of retinal diseases. The implications of these findings could extent to a broader scale. Indeed, the GPCR-mediated molecular machinery building up the phototransduction cascade is shared by other signal transduction processes, including chemotaxis, neurotransmission, cell communication, activation of olfaction and taste, and many others^49^. Understanding the mechanisms that influence this signaling cascade and achieving its controlled modulation is critical for new drug discovery, since about one-third of all drugs on the market target members of class A GPCRs^2^. More specifically, considering that GCAP1 is the major regulator of GC1 in human photoreceptors and that an increasing number of point mutations in its gene are associated with autosomal dominant COD or CORD, the development of novel biological therapies targeting this protein may help to restore the dysregulation of second messenger homeostasis in inherited retinal dystrophies, ultimately slowing or blocking cell death.

## Methods

### Cloning, expression, and purification of GCAP1 variants

Human myristoylated WT-GCAP1 was expressed in *E. coli* BL21 (DE3) after co-transformation with pBB131 containing the cDNA of *S. cerevisiae* N-myristoyl transferase (yNMT)^50^. The cDNA for E111V variant was obtained by PCR using QuikChange II Site-Directed Mutagenesis kit (Agilent) as described in ref. using the same expression protocol as for the WT. Proteins were purified according to the protocol elucidated in ref.^24^, briefly consisting of: (*i*) denaturation of inclusion bodies with 6 M Guanidine-HCl; (*ii*) refolding by dialysis against 20 mM Tris-HCl pH 7.5, 150 mM NaCl, 7.2 mM β-mercaptoethanol, and a combination of (*iii*) Size Exclusion Chromatography (SEC, HiPrep 26/60 Sephacryl S-200 HR, GE Healthcare) and (*iv*) Anionic Exchange Chromatography (AEC, HiPrep Q HP 16/10, GE Healthcare). The purity of GCAP1 variants was assessed by 15% acrylamide SDS-PAGE, samples were either exchanged against PBS, aliquoted and frozen with liquid nitrogen, or exchanged against ammonium bicarbonate, aliquoted and lyophilized. Protein samples were finally stored at −80°C.

The three-dimensional structure of human GCAP1 was obtained by homology modeling using the structure of Ca^2+^-loaded chicken GCAP1^51^ following the procedure illustrated in ref.^18^ *In silico* mutagenesis of E111V variant was obtained according to the protocol detailed in ref.^24^ The structures presented in **Fig. 1a** and **1b** were extracted from the last frame of 200 ns Molecular Dynamics simulations from ref.^24^, whose settings and protocols for energy minimization, equilibration and production phases were elucidated in refs^14,15^.

### Electrophoretic mobility shift assay

WT-GCAP1 and E111V-GCAP1 were dissolved in 20 mM Tris-HCl pH 7.5, 150 mM KCl, 1 mM DTT at a concentration of 30 μM, incubated for 5 min at 25°C with either 1 mM EGTA + 1.1 mM Mg^2+^ or 1 mM Mg^2+^ + 1 mM Ca^2+^, boiled, and run for 50 min at 200 V on a 15% acrylamide gel under denaturing conditions. Finally, protein bands were visualized by Coomassie blue staining.

### Circular dichroism spectroscopy

The effects of ion binding and of the E111V substitution on the secondary and tertiary structure of GCAP1 were evaluated by Circular Dichroism spectroscopy using a Jasco J-710 spectropolarimeter thermostated by a Peltier-type cell holder. Lyophilized proteins were dissolved in PBS pH 7.4 buffer at a concentration of 35 and 10 μM for near UV and far UV spectra, respectively. Five accumulations of each spectrum were recorded at 25°C in the absence of ions (500 μM EGTA for near UV, 300 μM for far UV) and after serial additions of 1 mM Mg^2+^ and Ca^2+^ (1 mM for near UV, 600 μM for far UV, leading to a free Ca^2+^ concentration of 500 and 300 μM, respectively). All spectra were subtracted with that of the buffer, near UV spectra were also zeroed by subtracting the average ellipticity between 310 and 320 nm, where no signal was expected.

### Dynamic Light Scattering

The hydrodynamic diameter of Ca^2+^-loaded WT-GCAP1 and E111V-GCAP1 was estimated by Dynamic Light Scattering using a Zetasizer Nano-S (Malvern Instruments, Malvern, UK). Proteins were dissolved in PBS pH 7.4 at 42 μM concentration and filtered with a Whatman Anotop 10 filter (20 nm cutoff, GE Healthcare) before starting the measurements. Samples were equilibrated for 2 min at 25°C and for each variant at least 100 measurements were collected, each consisting of 13 runs.

### Guanylate cyclase activity assay

GC1 enzymatic activity as a function of Ca^2+^ and GCAP1 concentration was measured after reconstituting WT-GCAP1 and E111V-GCAP1 with cell membranes of mGFP-GC1 cells (see below) previously extracted by lysis (10 mM HEPES pH 7.4, Protease Inhibitor Cocktail 1x, 1 mM DTT buffer) and 20 min centrifugation at 18000 x g, as previously described^30,52,53^. Cell membranes were resuspended in 50 mM HEPES pH 7.4, 50 mM KCl, 20 mM NaCl, 1 mM DTT and incubated with 5 μM GCAP1 variants at increasing [Ca^2+^] (<19 nM – 1 mM, controlled by Ca^2+^-EGTA buffer solutions^54^) to estimate the Ca^2+^ concentration at which cGMP synthesis by GC1 was half-maximal (IC_50_). To estimate the GCAP1 concentration at which GC1 activation was half-maximal (EC_50_), cell membranes were reconstituted with increasing amounts of each GCAP1 variant (0 - 20 μM) at low Ca^2+^ (<19 nM). Reported IC_50_ and EC_50_ values are represented as average ± standard deviation of 2 and 3 technical replicates, respectively. The statistical significance of the differences in IC_50_ and EC_50_ between WT-GCAP1 and E111V-GCAP1 was evaluated by means of two-tailed t-tests (p-value = 0.05).

### Conjugation of CF640R-N-hydroxysuccinimide (NHS) ester with WT-GCAP1

Far-red fluorescent dye CF640R (Biotium) was conjugated via NHS to WT-GCAP1 primary amines (mainly Lys residues, **Movie S1**) according to the manufacturer protocol. Briefly, GCAP1 was diluted in PBS pH 7.4 + 1 mM DTT to a final concentration of 76 μM in a final volume of 900 μl; then the solution was added with 100 μl sodium bicarbonate 1 M pH 8.3 and 2 CF640R-NHS aliquots previously resuspended in 50 μl total DMSO. The mixture was then wrapped in aluminum and incubated in rotation at RT for 1 h. Unconjugated dye was removed by washing 4 times the protein solution (see **Fig. S1b** for representative spectra of the 4 flowthrough) with PBS pH 7.4 for 10 min at 4400 x g and 4°C using an Amicon Ultra-4 concentrator with 3 kDa cutoff (Merck Millipore). The degree of labelling (DOL = 1.96) was calculated as the ratio between the concentration of dye in the protein solution measured based on the absorbance at 642 nm (ε= 105.000 cm^-1^ M^-1^), and the concentration of protein calculated by considering the dilution factor and the retention of Amicon concentrators (95%, according to manufacturer instructions). The concentration of free-CF640R in the protein solution was calculated by measuring the absorbance at 642 nm of wash 4, which was <1% with respect to protein concentration in all conjugation experiments. Unconjugated dye was blocked with 50 μl ethanolamine 1 M.

### Fluorescence spectroscopy

The emission fluorescence spectrum of 2 μM GCAP1^CF640R^ (645-680 nm) dissolved in PBS pH 7.4 was collected at 25°C on a Jasco FP-750 spectrofluorometer after excitation at 639 nm; the spectrum reported in **Fig. 1h** is an average of 3 accumulations after subtraction of the emission spectrum of the buffer in the same range.

### Liposome preparation

LPs were prepared by hydrating a thin lipid film of the same composition as photoreceptors rod outer segment membranes^55^ (phosphatidylethanolamine, phosphatidylcholine, phosphatidylserine, and cholesterol at a molar ratio of 40:40:15:5, Sigma) previously mixed in chloroform and dried in a speed-vac concentrator. Four mg of lipid film were hydrated with 1 ml PBS pH 7.4, vortexed for 30 min at room temperature, sonicated for 15 min in a water bath on ice and extruded 20 times through a 200 nm polycarbonate filter (Whatman). The encapsulation of CF640R, WT-GCAP1, E111V-GCAP1, or GCAP1^CF640R^ in LPs was achieved by dissolving the molecule to be loaded in PBS before lipid film hydration. Unencapsulated molecules were removed by washing at least 4 times the LPs suspensions with PBS pH 7.4 for 20 min at 4°C and 5000 x g using an Amicon Ultra-4 concentrator with 100 kDa cutoff (Merck Millipore). The degree of encapsulation was calculated by subtracting from the total mass of the molecule to be encapsulated that present in the flow-through and was found to be higher than 75% in all LP preparations.

### Nanoparticle Tracking Analysis

The concentration and size of LP suspensions were measured at 25°C by means of Nanoparticle Tracking Analysis on a NanoSight (Malvern) by recording 3 videos of 1 min each at 25 fps by setting 20 μl/min flow rate; camera level and detection threshold were automatically optimized for each measurement to maximize the signal-to-noise ratio. LP size reported in **Fig. 2** and LP concentration reported in **Table S1** represent the average ± standard error of 3 technical replicates.

### Fluorescence imaging of gel-immobilized liposomes

Stock suspensions of LPs, either filled with free-CF640R or empty, were diluted 1:400 v/v in 0.5% low gelling temperature agarose in Ames’ medium at 37°C (A1420; Merck). A thin film was polymerized over a pure agarose meniscus in a Petri dish and covered with Ames’ medium. 3D image stacks were acquired with a 63x/0.9NA water immersion objective and a CCD camera (DFC350 FX) in an upright widefield fluorescence microscope (DM LFSA) (all from Leica Microsystems) using a Cy5 filterset (49006; Chroma). Stacks were deconvolved and max projected along the z-axis using Fiji/ImageJ as detailed in ref.^56^.

### Generation of cGFP-GC1 and mGFP-GC1 stable HEK293 cell lines

HEK293 cells were cultured in DMEM medium supplemented with fetal bovine serum (10%, v/v), penicillin (100 units/ml) and streptomycin (100 μg/ml) at 37°C in humidified atmosphere with 5% CO_2_. Cells (6.25 x 10^5^) were seeded in 6-well plates in DMEM medium and grown overnight; the next day cell medium was replaced with OptiMEM reduced serum medium and cells were transfected using polyethyleneimine (PEI) as transfection reagent and 2 different vectors to obtain eGFP-expressing stable cell lines: (*i*) pIRES encoding for eGFP and human GC1 under different promoters, thus resulting in a cytosolic fluorescence (cGFP), and (*ii*) pcDNA3.1+N-eGFP encoding for GC1-eGFP fusion protein for localizing fluorescence on the membrane (mGFP). DNA (2.5 μg) was mixed dropwise to 10 μl PEI solution at a concentration of 1 μg/μl (DNA:PEI ratio of 1:5 w/w), added dropwise to 500 μl of pre-warmed OptiMEM, mixed and incubated 30 min at room temperature to allow DNA-PEI polyplex formation. Polyplexes were finally added dropwise to each well and the plate was incubated overnight at 37°C and 5% CO_2_. The next day, OptiMEM medium was replaced with DMEM and 48 h after transfection eGFP positive cells were selected using geneticin (500 μg/ml).

### Live-cell imaging

Cells (8 x 10^4^) were seeded in 4-well chambers (Ibidi) in DMEM medium; two days later the medium was replaced with OptiMEM reduced serum medium, then cells were incubated with 100 μl LP suspension per well (containing each ~ 0.4 mg lipid) and monitored in live-cell imaging. Experiments with fluorescently labelled GCAP1^CF640R^ were performed taking care of incubating the cells with the same nominal concentration of protein encapsulated in the LP aqueous core.

Live-cell imaging was performed using Leica TCS-SP5 Inverted Confocal Microscope equipped with temperature and CO_2_ controller and motorized stage that provides precise and automated acquisition of multiple fields of view. Images were collected simultaneously on different points of the sample immediately after cell-LP incubation and at 30 min interval for 24 h or 48 h total acquisition time. Images were captured after 488 nm and 633 nm laser excitation with a 40x objective (1.2 NA oil immersion) and further analyzed by Imaris 9.8 software (Oxford Instruments). The fluorescence intensity profiles of mGFP and LP-GCAP1^CF640R^ reported in **Fig. S4** were collected along the line across the cell shown in the insets using ImageJ.

### *Ex vivo* incubation experiments with mouse retinas

Experimental procedures on mice were performed in line with the permissions obtained by the Italian Ministry of Health (authorization n. 937/2021-PR referred to prot. n. 56DC9.74) and in accordance with the Italian (D.lgs.vo 116/92 and D.lgs 26/2014) and EU regulations (Council Directive 86/609/EEC).

Dark adapted adult C57Bl/6J mice were anesthetized with ketamine (80 mg/kg) + xylazine (5 mg/kg) and their retinas extracted through a corneal incision in room temperature Ames’ medium under dim red light. Animals were then immediately sacrificed with an overdose of anesthetic. After removing the vitreous each retina was placed in a plastic well containing 2 ml of incubation solution, and the wells inserted in an airtight box with a water layer at the bottom and a 95%O_2_/5%CO_2_ atmosphere. Incubation solutions consisted in the test suspension/solution diluted in Ames’ medium (from 1:17 to 1:66, depending on the experiment), taking care of reaching virtually the same final concentration for each suspension. The box was left floating in a water bath at 37°C. After the prescribed time the retinas were returned to room temperature Ames’ medium, made to adhere to black filter paper (AABP02500; Merck) with gentle suction and, optionally, sliced at 250 μm thickness with a manual tissue chopper. Image stacks were acquired as described for the imaging of gel-immobilized LPs, with 4x/0.1NA air, 20x/0.5NA and 40x/0.8NA water immersion objectives. Excitation was provided by an Hg lamp preheated to achieve stable output. Stacks were lightly deconvolved (Richardson-Lucy algorithm, 10 iterations) and average projected. Identical acquisition parameters were used when comparing retinas treated with different incubation solutions.

### Intravitreal injections

Mice were first anesthetized with ketamine (80 mg/kg) + xylazine (5 mg/kg), followed by application of eyedrops containing atropine and chloramphenicol (1%) + hydrocortisone (0.5%). Intravitreal injections were made under a stereomicroscope and dim blue light as follows: (*i*) a hole was made in the cornea near the *ora serrata* with the tip of a 31G insulin needle; (*ii*) glass micropipettes with a broken tip, connected to a 25 μl syringe (Hamilton) via PE tubing filled with mineral oil (330779; Merck), were front loaded with 2 μl of solution; (*iii*) the micropipette was inserted in the hole and the entire volume slowly injected in the vitreous. Mice were returned to their cages and allowed to recover in a paper blanket. After 20–24 h we performed retinal dissection, slicing and imaging as described for *ex vivo* incubations.

### Long duration *ex vivo* ERG recordings

Retina pairs were isolated as described for *ex vivo* incubations, made to adhere to white filter paper (SMWP02500; Merck) and placed at the bottom of plastic wells centered on a hole leading to the anode (**Fig. S5a**). In some experiments we dispensed with the filter paper and used instead small transparent cups to immobilize the retinas (**Fig. S5a**, inset labelled ‘variant’). Each well contained 2 ml of 40 μM AP4 (0101; Tocris) in sterile Ames’ medium and the cathode. Both electrodes were silver chloride wires inserted in an agar bridge. The wells were placed on an anodized aluminum platform covered with a layer of deionized water, inside a sealed incubation chamber purged with 95% O_2_/5% CO_2_. The temperature of the platform was actively controlled with a custom apparatus^57^. Small diameter PTFE tubing, leading from inside the wells to syringes residing outside the chamber, allowed injection and mixing of test solutions (100 μl) into the wells during the recordings with minimal perturbation. Immediately above the wells, attached to the lid of the chamber, an LED (505 nm; ND filters) delivered the same flash sequence every 15 min: (ph/μm^2^|no. of flash repetitions) 3.98|12, 8.27|10, 18.9|8, 50.5|6, 151|6, 510|4, 1660|3. Transretinal potentials were amplified by 5000, filtered in the band DC-100 Hz, digitized at 5 kHz and acquired with pClamp 9 (Molecular Devices). Electrophysiological records were analyzed in Axograph X with automated custom scripts. i_50_ was determined by fitting a Hill function to a plot of response amplitudes measured 90–130 ms after the flash (**Fig. S5b**). This range minimized the contribution of the very slow glial response and, in separate tests, gave parameter values close to the initial ones in BaCl_2_. TTP@i_50_ was estimated as the weighted average of the TTPs of the two flash responses straddling i_50_ (10 Hz Gaussian filtered records). Subsequent normalizations applied to these raw data are described in **Fig. S6** and legend. Treated and control retina positions in the wells were alternated from animal to animal to cancel out any environmental biases. In a limited number of tests (**Fig. 5h**) BaCl_2_ was injected in the wells using the syringe system after an initial stabilization period and stirred to obtain a final concentration of 50 μM.

## Data availability

The datasets generated during and/or analyzed during the current study are available from the corresponding author on reasonable request. Source data are provided with this paper.

## Acknowledgements

This study was supported by a grant from the Velux Stiftung (Project No. 1410) and by the PNRR Tuscany Health Ecosystem, milestone 8.9.1. The kind support of Retina Italia OdV is gratefully acknowledged. The Centro Piattaforme Tecnologiche of the University of Verona is acknowledged for providing access to spectroscopic and imaging platforms. We thank Carmen Longo for helpful discussions.

## Contributions

S.A. and L.C. performed the *ex vivo* and *in vivo* experiments. V.M., A.B. and G.D.C. performed *in vitro* experiments. A.A. performed *in cyto* and live imaging experiments. D.D.O. and L.C conceived and designed the study and supervised the project. All authors analyzed the data and contributed to design and write the manuscript with the supervision of L.C. and D.D.O. All authors have read and approved the manuscript.

## Supplementary information

This manuscript is published as a preprint together with a supplementary information file.

