## Supplementary materials for "Recombinant protein delivery enables modulation of the phototransduction cascade in mouse retina"

**This PDF file includes:**

Figures S1 to S6
Table S1
Captions for Movies S1 to S5

**Other Supplementary Materials for this manuscript include the following:**

Movies S1 to S5

**Figure S1**

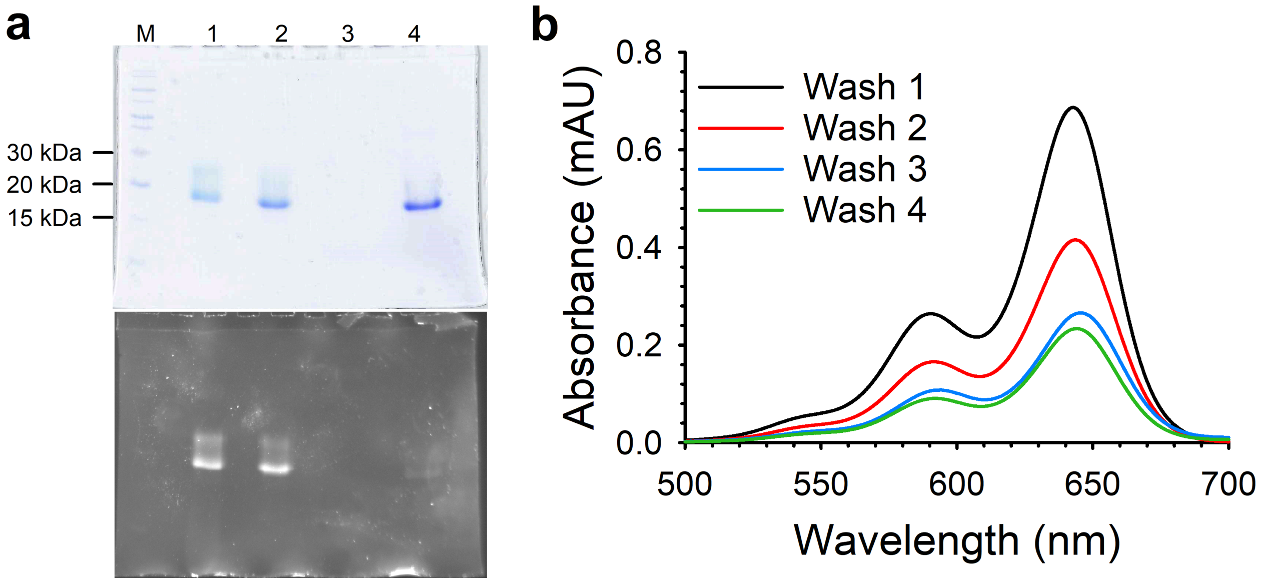


Assessment of the conjugation of GCAP1 with CF640R and removal of free dye. **a)** 15% SDS-PAGE of GCAP1 before and after conjugation with CF640R and after encapsulation in LPs. Lanes: M) marker, 1) LP-GCAP1$^{CF640R}$, 2) free-GCAP1$^{CF640R}$, 3) free-CF640R, 4) WT-GCAP1, stained with Coomassie blue (top panel) and upon excitation at 580 nm (bottom panel). **b)** Representative absorption spectra of 4 sequential washing steps to remove unconjugated dye.

#### Figure S2

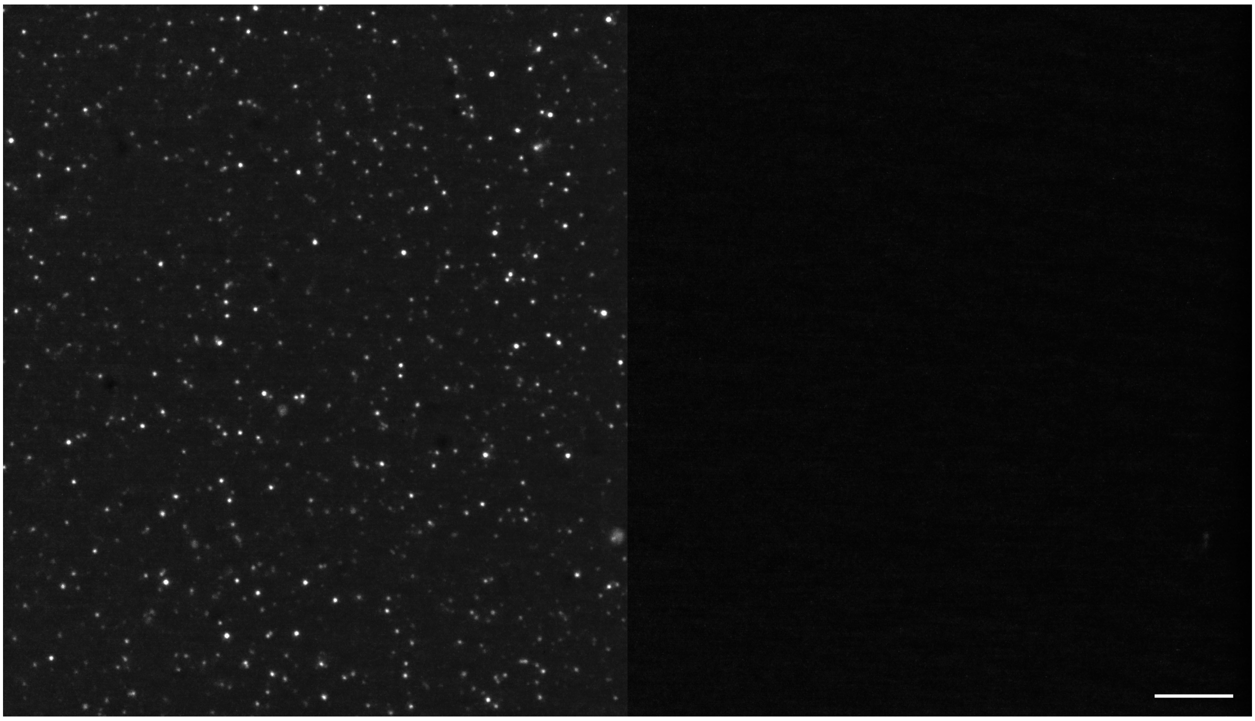

LP-CF640R appear as point-like (diffraction limited) fluorescence when suspended in agarose gel (left field) while empty LPs do not (right field). The two fields were acquired, processed, and displayed with identical parameters. Scale bar 10 µm.

#### Figure S3

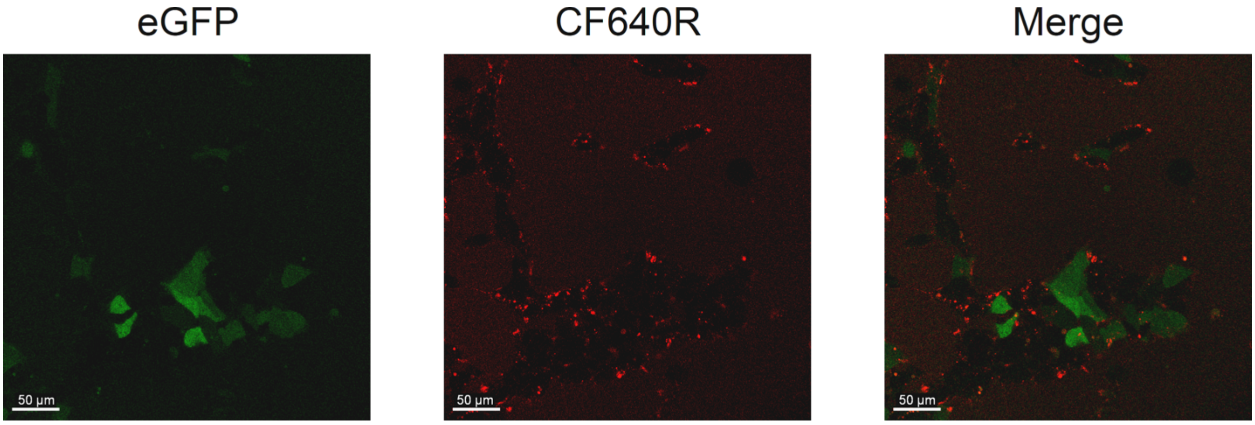


Representative images of cGFP cell line after 24 h incubation with 100 µl of 104 µM free-GCAP1$^{CF640R}$ after replacing cell medium with FluoroBrite DMEM. Left panel shows eGFP fluorescence, center panel shows CF640R fluorescence, left panel shows merged fluorescence.

### Figure S4

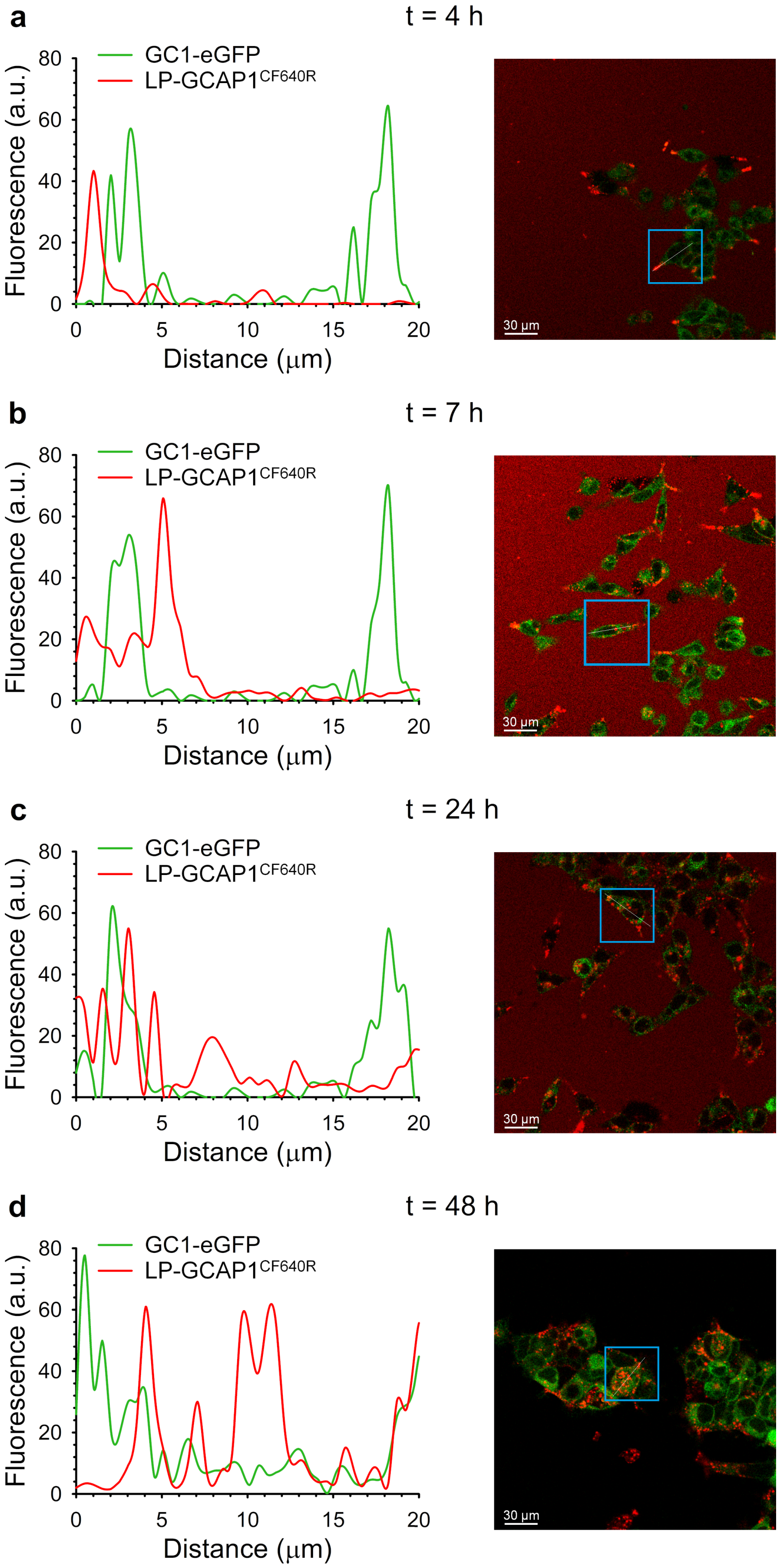
a
t = 4 h
b
t = 7 h
c
t = 24 h
d
t = 48 h
GC1-eGFP
LP-GCAP1CF640R
Fluorescence (a.u.)
Distance (μm)
0
20
40
60
80
0
5
10
15
20
30 μm

Representative (n= 6) fluorescence intensity profiles (left column) of live cell imaging (right column) of mGFP cell line (green) after **a)** 4 h, **b)** 7 h, **c)** 24 h and **d)** 48 h incubation with 100 µl of 4.6 nM LP-GCAP1$^{CF640R}$ (containing 27.4 µM GCAP1$^{CF640R}$ in the aqueous core, red). Fluorescence profiles were collected on the same z- plane as in **Fig. 3c**. Representative profiles refer to the cell framed in blue along the white line.

#### Figure S5

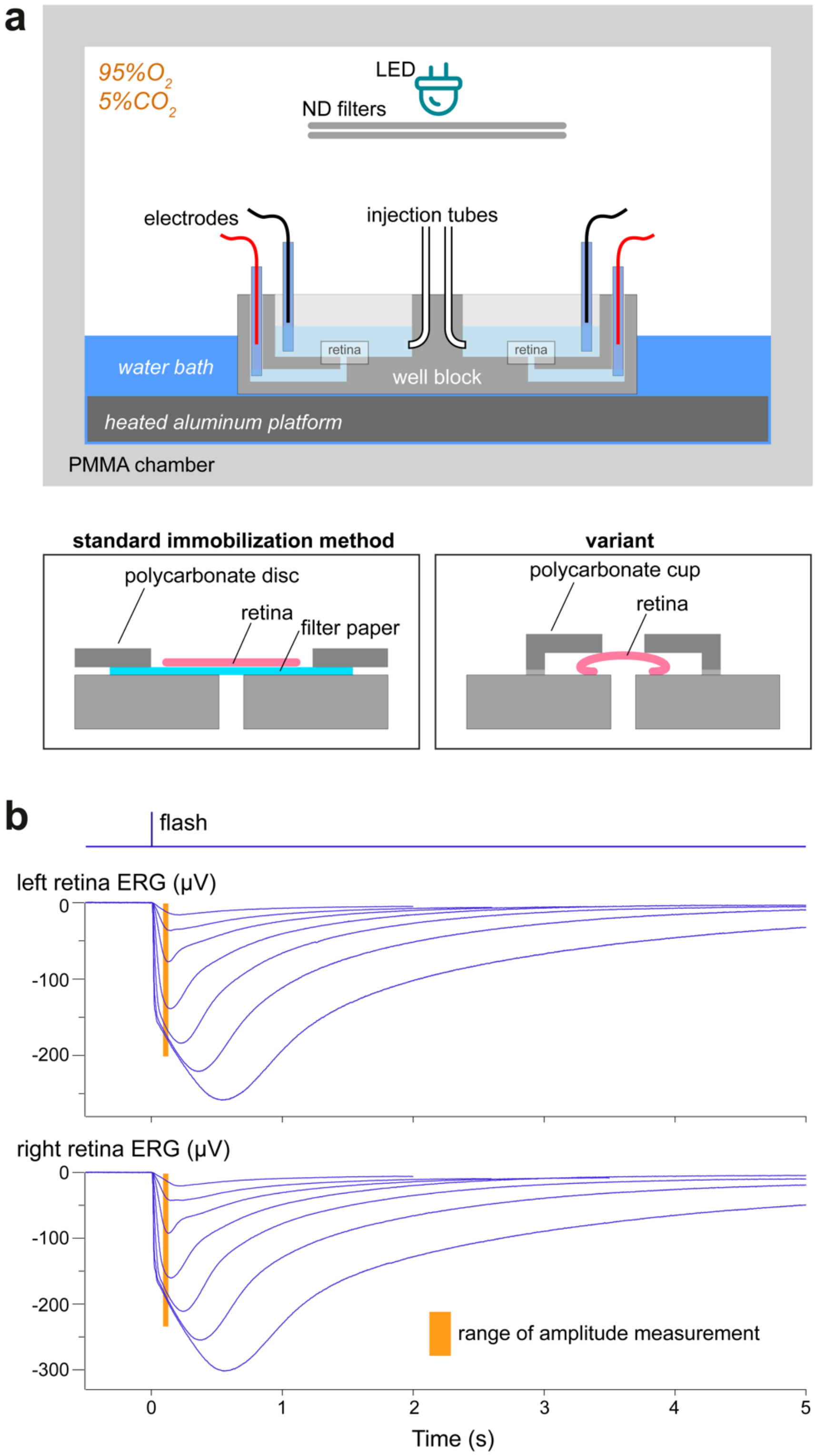


Custom designed *ex vivo* incubation and electroretinographic recording system. **a)** Schematic of the incubation and recording chamber (see Materials and Methods for details). The insets show two methods to immobilize the retina at the bottom of a well: on the left the standard approach of using a filter paper support, while on the right the retina is held slightly compressed by a small polycarbonate cup. This latter method is mechanically less stable but avoids any capture of LPs or free proteins by the porous filter paper. **b)** Examples of concurrently recorded flash responses from the two eyes of a mouse. Each trace is the average of several flash responses, with an entire flash family being delivered every 15 min. Flash strengths (ph/µm$^2$)|no. of repetitions: 3.98|12, 8.27|10, 18.9|8, 50.5|6, 151|6, 510|4, 1660|3. Light sensitivity ($i_{50}$) was estimated by fitting a Hill function to the response amplitudes measured in the range 90–130 ms after the flash (orange areas). Kinetics (TTP@$i_{50}$) was estimated as the time to peak of the hypothetical response at $i_{50}$. Left retina: $i_{50}$ = 23.2 ph/µm$^2$; TTP@$i_{50}$ = 136 ms. Right retina: $i_{50}$ = 20.7 ph/µm$^2$; TTP@$i_{50}$ = 136 ms.

#### Figure S6

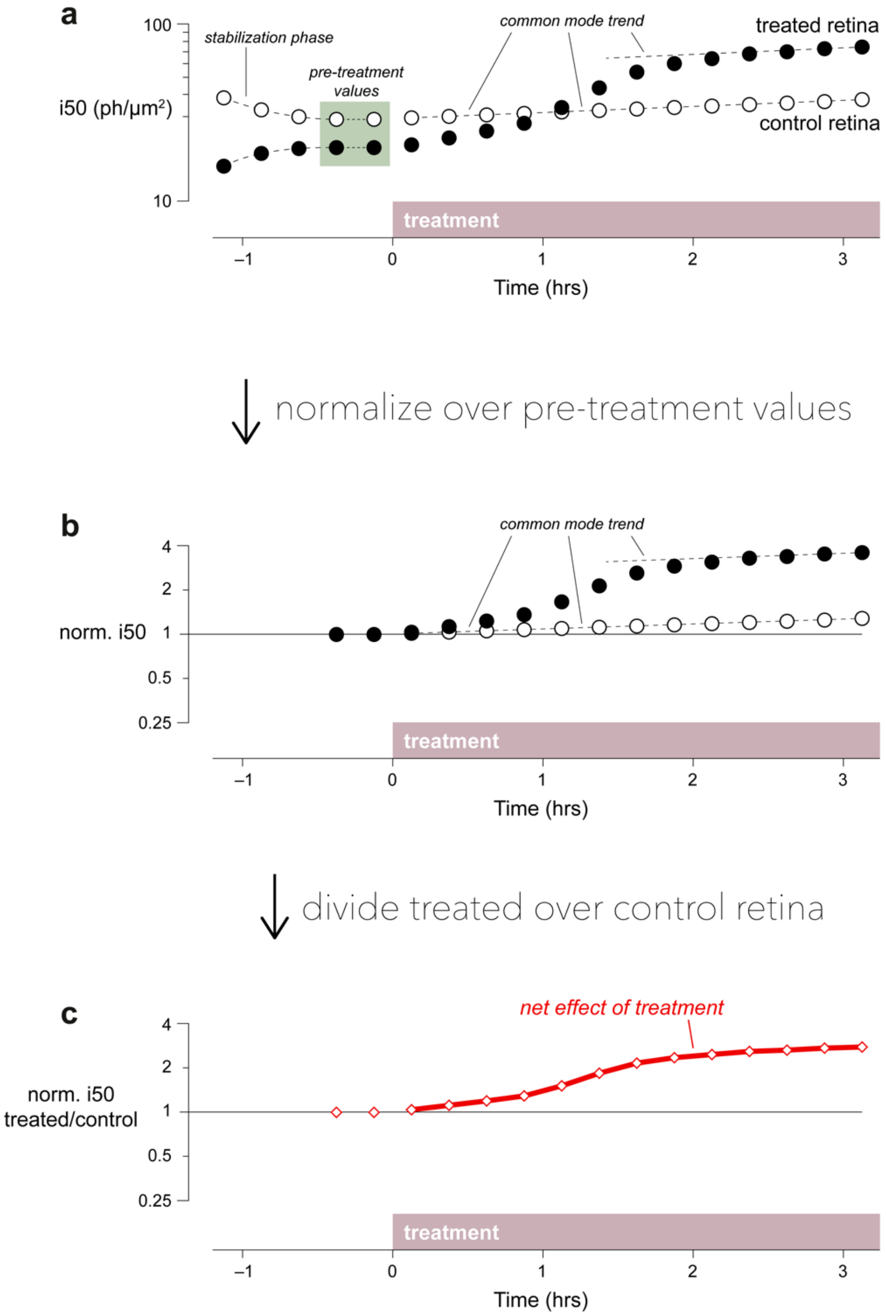


Processing pipeline of the *ex vivo* ERG data ($i_{50}$ or TTP@$i_{50}$) obtained in a hypothetical experiment where a retina is incubated with a test drug, while the other retina is given vehicle (i.e. control). **a**) We assume that the two retinas, being from the same animal, behave identically except for (*i*) an initial stabilization phase due to differences in their isolation and manipulation, and (*ii*) a scaling factor in their steady state light sensitivity due to their geometry in the incubation and recording chamber. **b**) We proceed to normalize the two sets of raw values over their respective pre-treatment levels. **c**) We remove any trends that are common to both retinas by dividing the normalized values of the treated retina by those of the control. We are left with a single time series that shows the net effect of the tested drug: in the example an over twofold decrease in scotopic light sensitivity.

#### Table S1

| Liposomes | Size (nm) | Concentration (nM) |
|---|---|---|
| LP-empty | 158.6 ± 1.1 | 5.1 nM |
| LP-E111V-GCAP1 | 153.3 ± 2.9 | 4.3 nM |
| LP-GCAP1$^{CF640R}$ | 168.7 ± 0.7 | 4.6 nM |
| LP-empty (day 1) | 149.1 ± 3.0 | 4.6 nM |
| LP-empty (day 90) | 151.0 ± 5.4 | 3.2 nM |
| LP-empty (day 180) | 157.4 ± 2.0 | 4.0 nM |
| LP-E111V-GCAP1 (day 1) | 164.3 ± 0.4 | 2.9 nM |
| LP-E111V-GCAP1 (day 90) | 152.1 ± 3.4 | 2.6 nM |
| LP-E111V-GCAP1 (day 180) | 153.7 ± 1.8 | 3.0 nM |
| LP-CF640R | 160.5 ± 1.2 | 3.9 nM |

Size, concentration, and stability over 180 days of LPs loaded with different molecules (dissolved in PBS) monitored by Nanoparticle Tracking Analysis. Data refer to the mean ± standard error of 3 technical replicates.

#### Movie S1

The three-dimensional structure of GCAP1 is shown as light-grey cartoon with the molecular surface in transparency, $Ca^{2+}$-ions are displayed as green spheres, the sidechains of Lys residues are labelled represented as red sticks with N atoms in blue. The molecular surface of the primary amines belonging to Lys sidechains is shown in blue in transparency.

#### Movie S2

Live-cell imaging at 6 h of cGFP cell line incubated with 100 µl of 140 µM free-CF640R, snapshots were acquired at a 30 min interval, green fluorescence refers to eGFP, red fluorescence refers to free-CF640R molecules.

#### Movie S3

Live-cell imaging at 6 h of mGFP cell line incubated with 100 µl of 104 µM free-$GCAP1^{CF640R}$, snapshots were acquired at a 30 min interval, green fluorescence refers to eGFP, red fluorescence refers to free-$GCAP1^{CF640R}$ molecules.

#### Movie S4

Live-cell imaging at 6 h of cGFP cell line incubated with 100 µl of 104 µM free-$GCAP1^{CF640R}$, snapshots were acquired at a 30 min interval, green fluorescence refers to eGFP, red fluorescence refers to free-$GCAP1^{CF640R}$ molecules.

#### Movie S5

Live-cell imaging at 24 h of mGFP cell line incubated with 100 µl of 4.3 nM LP-$GCAP1^{CF640R}$ (containing in the aqueous core the equivalent number of $GCAP1^{CF640R}$ molecules present in a 104 µM solution) snapshots were acquired at a 30 min interval, green fluorescence refers to eGFP, red fluorescence refers to LP-$GCAP1^{CF640R}$.
